# Longitudinal Subject Pairing in Cross-Sectional Neonatal Data Reveals Asynchronous Structural and Functional Brain Maturation

**DOI:** 10.64898/2026.07.08.737182

**Authors:** Roxana Namiranian, Mohadeseh Sadeghi, Hamid Abrishami Moghaddam

**Author notes:** Senior author (Hamid Abrishami Moghaddam).

## Abstract

The asynchronous development of structural and functional brain networks in early childhood remains largely unexamined, primarily due to the scarcity of longitudinal neuroimaging data. Resolving this temporal dimension is critical, as it promises to reshape our understanding of structural-functional (S-F) coupling, revealing not only whether brain architecture supports function, but also when and over what timescale its influence emerges. However, while rapid neonatal brain maturation and logistical constraints continue to hinder longitudinal data collection, large-scale cross-sectional multimodal datasets are currently available to bridge this gap. Here, we propose a longitudinal subject-pairing framework that reconstructs developmental trajectories from cross-sectional data. It pairs the infants with a predefined age gap while maximizing their similarity in both structural and functional features, thereby approximating the longitudinal trajectory of functional changes in relation to structural maturation. As a case study, we applied this framework to the perisylvian region in a subset of 505 neonates from the multimodal dHCP brain dataset. The myelination index was derived as a structural feature from MRI, and the fractional amplitude of low-frequency fluctuations (fALFF) was derived as a functional feature from resting-state functional MRI. A conventional cross-sectional analysis revealed a moderate S-F correlation magnitude (r = 0.34). In contrast, the proposed framework demonstrated a significant increase in S-F coupling to r = 0.46 when the structural maturation precedes functional maturation by approximately five days. These findings provide novel evidence of a functional maturation lag relative to structural brain development in neonates. Beyond elucidating S-F relationships in the early developing brain, this work establishes a framework for future longitudinal studies and advances in brain modeling across developmental trajectories, aging, and disease prediction.

## INTRODUCTION

Understanding the mechanisms linking structural maturation to functional reorganization in the developing human brain remains a fundamental challenge in neuroscience. Specifically, the early postnatal period, characterized by rapid and profound neurodevelopment, provides a critical window for observing this interplay, thereby revealing the principles of structural-functional (S-F) co-development^1–4^. However, investigating this relationship during early development presents challenges, as several studies report weak or insignificant S-F couplings^5–10^. Proposed explanations for these findings include the use of analytical features ill-suited to the developing brain^6,7,11,12^, and the heterogeneous, regionally distinct maturation trajectories of S-F coupling^5,7,10,13,14^.

For a long time, research into this S-F relationship across early development has been governed by the prevailing paradigm that structural and functional maturation progress hand-in-hand^3^. Yet, a pivotal question has received little attention: do structural and functional maturation unfold concurrently, or does a synchrony assumption─arising from methodological limitations─obscure a systematic chronological relationship?^8,12,14^ Owing to the logistical constraints of acquiring large-scale multimodal longitudinal brain data^12,15–18^, most studies continue to examine early structural and functional development as temporally static phenomena, implicitly assuming that their trajectories evolve in tandem^3,7,8,12,14,19–21^. Consequently, the possibility that structural maturation serves as a foundation for brain function, potentially exerting delayed effects on functional development, has remained largely unexplored. This limitation is compounded by a lack of analytical frameworks capable of quantifying such temporal offsets, thereby hindering efforts to robustly test the temporal dynamics of S-F relationships. Resolving this temporal dimension is critical, as it promises to reshape our understanding of S-F coupling, revealing not only whether brain architecture supports function, but also when and over what timescale its influence emerges.

Longitudinal designs represent the conventional methodological choice for investigating the temporal dynamics of S-F relationships. Although robust for tracking developmental changes, this approach is associated with significant practical challenges, including the substantial effort required for data acquisition across frequent imaging sessions, a difficulty particularly amplified in neonatal populations where cerebral development is exceptionally rapid^1,2^. Furthermore, the inherent complexity of neurodevelopment necessitates multiple large-scale multimodal longitudinal cohorts to fully elucidate the temporal dynamics governing asynchronous S-F interactions. Of note, major initiatives have focused on assembling large-scale cross-sectional datasets, such as the developing Human Connectome Project (dHCP), providing an invaluable foundational resource for the early brain connectome. While these datasets offer precise gestational age information, their potential to interrogate temporal dynamics and precedence in S-F maturation has remained largely untapped.

Our work addresses this gap by introducing a novel methodological framework that repurposes crosssectional data to infer developmental trajectories, thereby circumventing the limitations of longitudinal studies while extracting novel insights from existing resources. Our algorithm identifies pairs of distinct infants within the cohort separated by a specified age difference to simulate a developmental sequence between structure and function. It links neonates across developmental stages on the basis of the distance between their respective structural and functional profiles. The new algorithm belongs to a broader class of trajectory-reconstruction approaches that infer temporal dynamics from cross-sectional observations when direct longitudinal tracking is impractical or impossible^22,23^. As a case study, we demonstrate that hypotheses regarding the delayed influence of brain structure on function can be tested using longitudinally paired cross-sectional data, without requiring strictly longitudinal observations. Specifically, we test hypotheses regarding the scale of this temporal delay, demonstrating that the influence of structural changes on function operates on the order of days.

## METHODS

### Data and Participants

This study utilized a subset of the second-release dataset from the dHCP^24^, which comprises multi-modal neuroimaging data from 505 neonates. In line with our previous work^11^, the current analysis was restricted to term-born neonates (gestational age (GA) ≥ 37 weeks). Following the exclusion of subjects exhibiting excessive head motion during fMRI acquisition^25^ or those with incomplete data, the final cohort comprised N=166 imaging sessions (81 female; mean GA at scan ± STD: 41.3 ± 1.6 weeks; range: 38-44.7 weeks). The age distribution of the neonatal cohort, stratified by gender, is provided in Supplementary **Figure S1**. All infants were scanned during natural sleep using a 3T Philips Achieva scanner equipped with a dedicated neonatal head coil at the Evelina Neonatal Imaging Centre, London.

The structural imaging protocol consisted of acquiring high-resolution T1-weighted and T2-weighted images. The final reconstructed structural data, achieved through slice-to-volume registration, had an isotropic resolution of 0.5 mm^26^. Resting-state fMRI (rs-fMRI) was acquired over a 15-minute period using a multiband Echo-Planar Imaging (EPI) sequence (2.15 mm isotropic, TR=392 ms). Data preprocessing was performed via the standard dHCP pipeline, which included distortion correction, motion correction, registration to structural space, high-pass filtering, and ICA-based denoising^27,28^.

### Parcellation and Feature Extraction

Neonatal brains were parcellated into 90 distinct anatomical regions using the University of North Carolina (UNC) infant atlas^29^, which provides a fine-grained regional division around the Sylvian fissure. The current analysis focused specifically on 16 perisylvian regions (PSRs) per hemisphere, including the superior temporal gyrus (STG), middle temporal gyrus (MTG), Heschl’s gyrus (HG), supramarginal gyrus, angular gyrus, and the insula. To align the UNC atlas with individual participant data, age-specific templates were registered to each neonate’s native T2-weighted (T2w) structural space using both linear and nonlinear transformations. For functional analysis, these volumetric masks were subsequently transformed into native functional space utilizing the transformation matrices provided by the dHCP consortium. For structural analysis, the parcels were mapped to the cortical surface using the HCP Connectome Workbench.

In accordance with recent findings^8,11^, the fractional amplitude of low-frequency fluctuations (fALFF) was selected as the functional feature of interest due to its established correlation with the myelination index, which served as the microstructural feature. The fALFF metric was computed for each perisylvian subregion directly from the preprocessed blood-oxygenation-level dependent (BOLD) time series. Concurrently, the myelination index, quantitatively defined as the T1-weighted to T2-weighted (T1w/T2w) intensity ratio, was derived from the surface-based maps generated by the standard dHCP pipeline^26^. To obtain regional metrics, both fALFF values and T1w/T2w ratios were subsequently averaged across all vertices within each respective UNC parcel.

To ensure that the observed structure-function relationships were not driven by common dependencies on maturational factors, we statistically controlled for postmenstrual age (PMA) at scan. Prior to the main analytical steps, the effect of this confounding variable was regressed out from both the fALFF and T1w/T2w metrics, and the resulting residuals were used in all subsequent analyses. Here, the S–F coupling was determined by selecting the maximum correlation coefficient within the correlation matrix derived from the fALFF and myelination feature vectors.

### Longitudinal Pairing Algorithm for Inferring Developmental Precedence

To rigorously investigate the developmental precedence between functional and structural maturation in the absence of conventional longitudinal data, we developed a novel longitudinal pairing framework. This algorithm facilitates the construction of synthetic developmental trajectories by systematically matching and integrating data across distinct individuals within the cohort. A schematic overview of this subject-pairing algorithm is illustrated in **Figure 1**.

**Figure 1.**
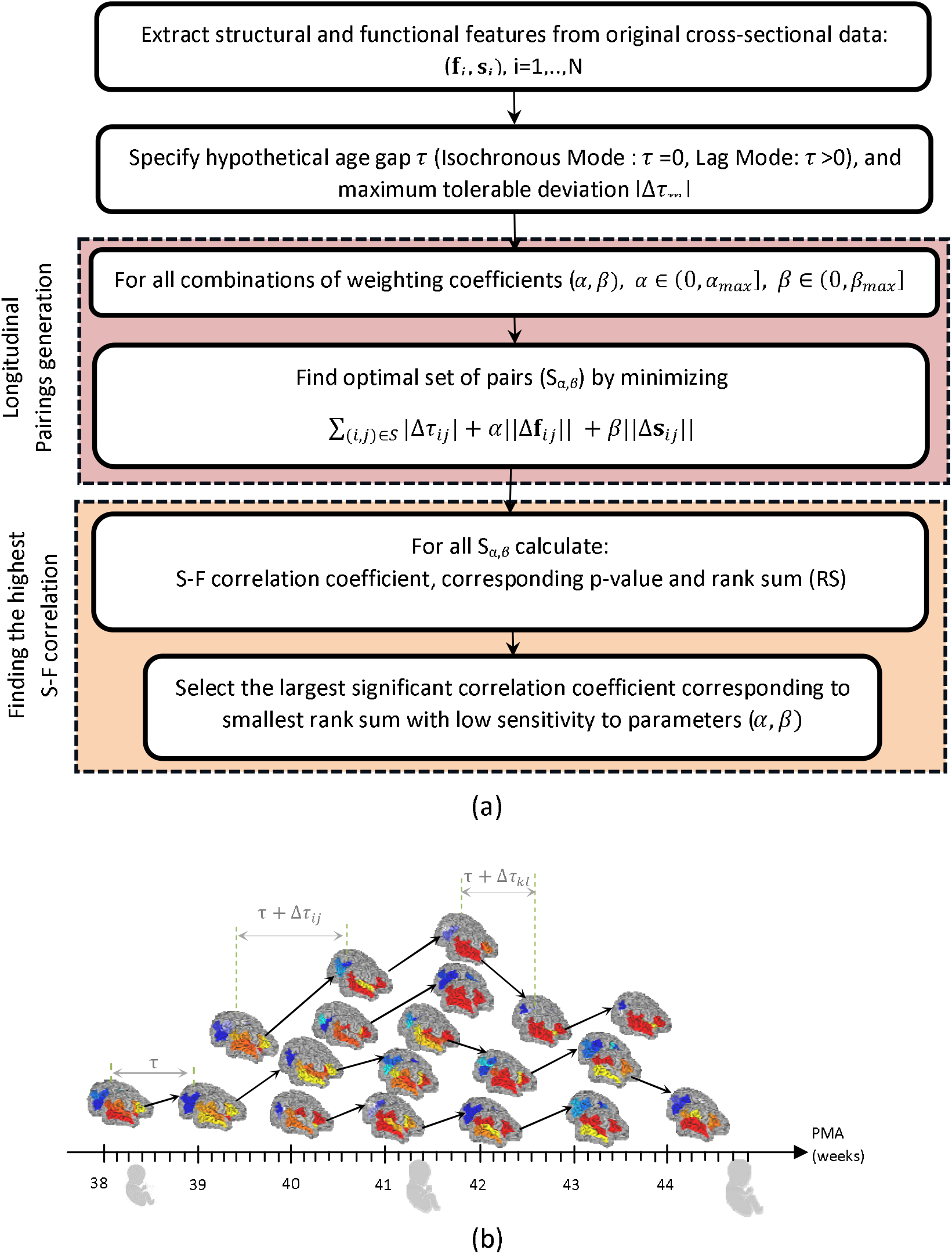
An overview of the proposed framework to study asynchronous structural-functional neurodevelopment. (a) The longitudinal subject-pairing algorithm to study developmental precedence using crosssectional data. (b) Schematic representation of the subject-pairing algorithm. Each arrow represents a pairing between the structural feature (red colormap) of a given subject and the functional feature (blue colormap) of another subject, defined by a target age lag of r = 7and |Δr_m_|=2 days. Here, Δr_ij_ (Δr_kl_) indicates the tolerated deviation around the specified age gap r between the ith (lth) and jth (kth) subjects.

Our algorithm identifies pairs of infants separated by a specified age difference (r) to simulate a developmental sequence. To accommodate the discrete nature of the gestational age sampling within the dHCP dataset, we defined a tolerance window [-|Δr_*m*_|,|Δr_*m*_|I around the target age difference r, effectively accepting any age gap within the interval [r-|Δr_*m*_|,r + |Δr_*m*_|I. For a given target gap r, we first identified all candidate pairs whose age difference fell within this defined tolerance. Each candidate pair (*i*,*j*) was subsequently scored using a cost function that penalizes both deviations from the hypothetical age gap and respective dissimilarities between the subjects’ functional and structural profiles:

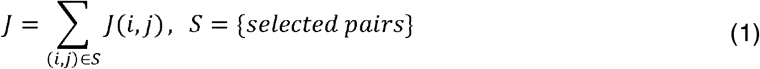

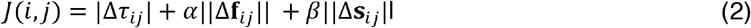

Here, Δ*r*_*ij*_ is the tolerated deviation from the hypothetical age gap r. The terms ||Δf_*ij*_ || and ||Δs_*ij*_ || denote the Euclidean distances between the functional and structural feature vectors of the two infants, respectively. The coefficients α and β are weighting parameters that balance the relative contributions of the age gap deviation, functional dissimilarity, and structural dissimilarity, respectively. By minimizing this objective function, the algorithm effectively pairs subjects with highly congruent structural and functional profiles. To intuitively illustrate the interplay of these terms, consider—without loss of generality—the case of pairing subjects of identical age to exchange their structural and functional features. In this scenario, the optimal candidates are those who share the exact same chronological age and exhibit the lowest cross-subject variability in features, a constraint enforced by the final two distance terms. However, a subject of a slightly different age (e.g., one day older or younger) may exhibit greater feature similarity to a target subject than an exact age-match. This demonstrates the critical regularizing role of the first term, which permits controlled age flexibility to maximize phenotypic matching.

Consistent with our methodological objective—namely, leveraging cross-sectional data to derive synthetic longitudinal trajectories—we aim to pair subjects across a target age gap *r*. Minimizing the objective function ensures that while the selected subject pairs maintain a target age separation (*r*), they are explicitly constrained to minimize baseline cross-subject structural and functional discrepancy. Consequently, the resulting feature differences within each pair can be more reliably attributed to true developmental progression rather than confounding inter-individual variability.

The algorithm initiates by evaluating the cost function for all eligible subject pairs. These pairs are subsequently ranked in ascending order of cost, and the optimal selections are determined via a greedy optimization approach that strictly enforces a feature exclusivity constraint, ensuring that each individual subject’s data is uniquely assigned only once. For the final analysis, data from the selected optimal pairs are fused based on one of two specified age gap configurations:

- **Lag Mode**: Within each paired unit, the structural data acquired from the chronologically younger subject are correlated with the functional data from the older subject.
- **Isochronous (Zero-Lag) Mode**: Pairs exhibiting a target age gap of zero (*r*= 0) are integrated to evaluate contemporaneous relationships, assuming no developmental precedence between the structural and functional data streams.

Within the theoretical context of an ideal isochronous analytical mode, an individual subject’s structural data are conceptually paired directly with their concurrently acquired functional data. This self-pairing configuration (s_*i*_ →.f_*i*_) constitutes the standard methodology employed in traditional cross-sectional investigations, where the structural features of a given participant are correlated against their corresponding, co-measured functional metrics. Conversely, the novel pairing framework introduced in this study departs fundamentally from this conventional paradigm by explicitly prohibiting self-pairing within the isochronous control mode. This deliberate methodological constraint is strictly required to preserve internal consistency and ensure robust statistical comparability between the results derived from the novel lag-based mode and the baseline isochronous control condition.

### Exhaustive search for the highest significant S-F correlation

Minimization of the objective function (1), with different values of weighting coefficients (α, β), results in distinct combinations of paired subjects. Obviously, the appropriate selection of these coefficients influences the quality of the resulting set of paired subjects. To identify the optimal values for α and β, we employed an exhaustive search strategy within a multi-objective optimization framework. The parameter space was nonuniformly discretized with sufficient precision within a predefined range of values. Within this discrete space, we looked for combinations of paired subjects that simultaneously satisfy the following criteria:

- First, these sets yield the maximum attainable and significant S-F correlation values. To evaluate the statistical significance of the S-F correlations derived from the optimal subject pairs under each parameter configuration, permutation testing was performed. For each age gap (*r*) and parameter set (α, β), an empirical null distribution was constructed by shuffling the pairings between structural and functional data across the entire eligible cohort. Specifically, structural and functional metrics were randomly reassigned to generate new pairings from the infant pool, and the S-F correlation was recomputed across 500 independent permutations. The p-value was defined as the proportion of permuted datasets yielding a Fisher’s z-transformed correlation equal to or greater than the observed correlation derived from the optimized pairs. Correlations surviving a significance threshold of p < 0.05 were deemed statistically significant and unlikely to have occurred by chance.
- Second, the average absolute deviation from the target age difference, as well as the average distances of structural and functional features between paired subjects, are minimized. Notably, because these three metrics are expressed on different scales, we utilized their ranked scores rather than their raw values to facilitate simultaneous minimization. Accordingly, we sought a set of weighting parameters (α, β) that yields a set of paired subjects that simultaneously maximizes the S-F correlation and minimizes the rank sum (RS) score:

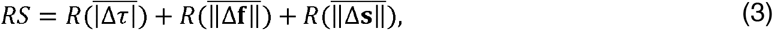

Where 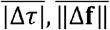, and 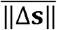 denote the average values of |Δr_*ij*_|, ‖Δf_*ij*_‖, and ‖Δs_*ij*_‖, respectively, computed across the selected pairs corresponding to coefficient sets *(α*, *β)*, and 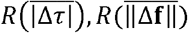, and 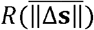 represent their respective ranks.
- Finally, the resulting correlation values and rank sums exhibit low sensitivity to perturbations in the weighting parameters. In other words, the optimal correlation and rank sum values vary only marginally across a broad range of weight changes.

### Comparison with Baseline Correlation Values

To evaluate the performance of our longitudinal subject-pairing algorithm, we applied it to gather further evidence of lagged functional brain development in neonates. Accordingly, the S-F correlation values resulting from our pairing algorithm were compared to those derived from (1) original cross-sectional data and (2) our algorithm operating in isochronous mode (r =0). The former serves as an ideal crosscorrelation baseline under the hypothesis of concurrent structural and functional development, whereas the latter provides a negative control value representing synchronous S-F development realized by synthetic pairs with zero age gap.

It is worth noting that the number of unique infant pairs under either lag or isochronous modes may fall slightly below the original cohort size. The underlying rationale for this constraint is illustrated in the schematic overview of the pairing algorithm in **Figure 1(b)**. As shown, when matching *n*_*i*_ subjects at age *t*_*i*_with *n*_*i*_ available subjects within the age range[t_*i*_+r - |Δr_*m*_|,t_*i*_+r + |Δr_*m*_|], the number of possible matches is limited to *min*(*n*_*i*_, *n*_*j*_). Consequently, when *n*_*j*_ < *n*_*i*_, no corresponding pair exists for certain subjects at age *t*_*i*_.

To ensure that comparisons of S-F correlations were not confounded by variations in sample size or age distribution, we implemented a subsampling procedure for the original dataset. Specifically, when estimating the S-F correlation for individuals in cross-sectional data, we randomly resampled a subset of infants equal in size to the number of paired subjects generated for a given mode, while explicitly matching the age distribution of the original data. This procedure was repeated iteratively to generate a baseline distribution of S-F correlation estimates under the conventional cross-sectional approach, thereby enabling an unbiased comparison with our pairing algorithm.

Additionally, to ensure a fair comparison across different lag conditions, the number of pairs in the isochronous mode (*r* = 0) was adjusted to closely match the number of pairs available in each lag mode. If the number of pairs identified in a lag mode is smaller than the number obtained through an exhaustive search in isochronous mode, the latter are ranked according to their selection frequency across different values of the weighting coefficients. The most frequently selected pairs are then used to compute the S–F correlation. Conversely, if the number of pairs identified in a lag mode exceeds that in the isochronous mode, a subset of lag-mode pairs equal in size to the isochronous pairs can be selected to ensure a balanced comparison. In the present study, however, the maximum number of available isochronous pairs was retained because the lag-mode pairs exceeded the isochronous mode marginally and only for a single lag condition. This procedure allowed the observed differences in correlation strength to be primarily attributed to the temporal delay itself, rather than to variations in sample size. Accordingly, the *r* = 0 condition served as an additional reference, enabling the specific effect of temporal delay to be isolated while minimizing potential biases arising from differences in the pairing procedure.

## RESULTS

A maximum deviation from the specified age difference |Δ *r* _*m*_|=2 days was considered in both lag and isochronous modes. While a maximum deviation of one day would align with the precision of neonatal age measurements in the dHCP, it might considerably reduce the number of feasible pairings, thereby excessively restricting the search space. Therefore, a ±2-day tolerance for the specified age gap was adopted as a compromise between temporal precision and the number of pairable subjects within the search space. The maximum values for the weighting coefficients, α_*max*_ and β_*max*_, were set to 50 in both the isochronous and lag modes. **Table S1** and **S2-S11** in the supplementary materials indicate the optimal pairing results for each combination of weighting coefficients in isochronous (*r* =0) and lag modes (*r* = 1, …,10 days), respectively. In these tables, the first two columns list the evaluated weighting coefficients. Each row provides the number of optimal pairs along with the corresponding S-F correlation and rank sum values. **Figure 2(a-c)** depict the correlation map, rank-sum values, and their corresponding ratio, respectively, within the weighting coefficient space. These values were calculated using optimal subject pairs generated from non-uniformly discretized coefficients at a lag mode of *r*= 5 days. As expected, the correlation and rank sum values of the optimal pairs remain largely invariant along the line defined by the slope. This consistency occurs because the primary role of the weighting coefficients, α and β, is to normalize the different terms on the right side of the cost function (1) to the same scale. Consequently, the optimal pairs generated by identical β/α ratios in (1) are substantially similar. The contoured bright region in **Figure 2(c)** highlights an extensive domain where the weighting coefficient pairs yield the highest correlation to rank-sum ratios. **Figure S2** in the supplementary materials illustrates similar maps for isochronous (*r* =0) and lag modes (*r* =1-10 days).

**Figure 2.**
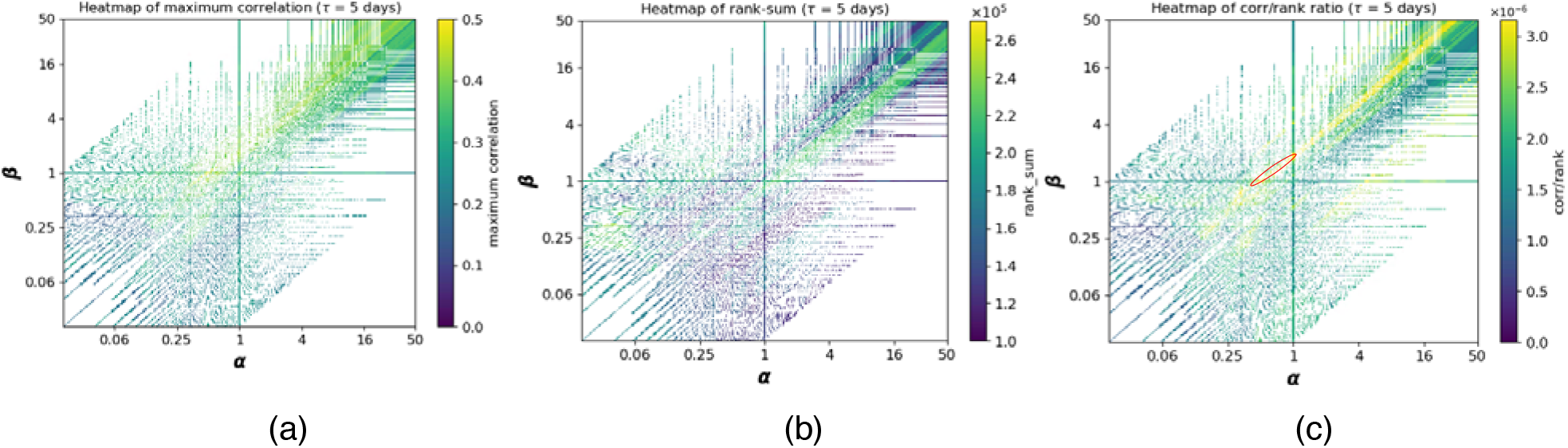
The heatmap of the S-F correlation and rank sum values computed on the optimal set of pairs corresponding to lag mode(r =5 days) in the nonuniformly discretized coefficients space. (a) S-F correlation coefficient, (b) rank sum value, and (c) their ratio. The S-F correlation and rank sum values of the optimal pairs remain mostly unchanged along the line defined by the slope β/α.

Figure 3. shows the final results achieved by the exhaustive search in the parameter space to identify the highest significant S-F correlation. The red circles represent the S-F correlation values corresponding to each age gap in lag mode, obtained by maximizing the correlation while minimizing rank sum and ensuring low sensitivity to parameter variations. **Table S12** in the supplementary materials indicates how this procedure was implemented to determine the S-F correlation values. Furthermore, **Figure 3** compares the S-F correlation values computed in lag mode against those derived from two distinct baselines: (1) individuals from the original cross-sectional dataset, and (2) paired subjects determined by our algorithm operating in isochronous mode (*r* =0). Accordingly, the boxplots depict the distribution of SF correlation values derived from the original cross-sectional dataset after resampling, with sample sizes matched to the number of optimal pairs (*n*) associated with each age gap in lag mode. This distribution represents the baseline cross-correlation under the hypothesis of concurrent structural and functional development. Finally, the blue squares denote the S-F correlations computed in isochronous mode, where the number of pairs (*n*) closely matches the number of available pairs for each age gap in lag mode. These data points serve as negative controls, representing synchronous S-F development as realized by synthetic pairs with zero age gap computed by our algorithm.

Higher S-F correlations are expected when computed for individuals within a cross-sectional cohort compared to synthetic pairs generated by our longitudinal pairing framework in isochronous mode, because the coupling between the structural and functional features within the same individual is inherently stronger than the correlation between a structural feature from one subject and a functional feature from a distinct, age-matched subject. Especially, we intentionally inhibit the algorithm from pairing the structural and functional features of the same subject. Consequently, the S-F correlations obtained in lag mode must be benchmarked against the baseline correlations computed in isochronous mode rather than against the intra-subject S-F correlation derived from the baseline cross-sectional dataset.

**Figure 3.**
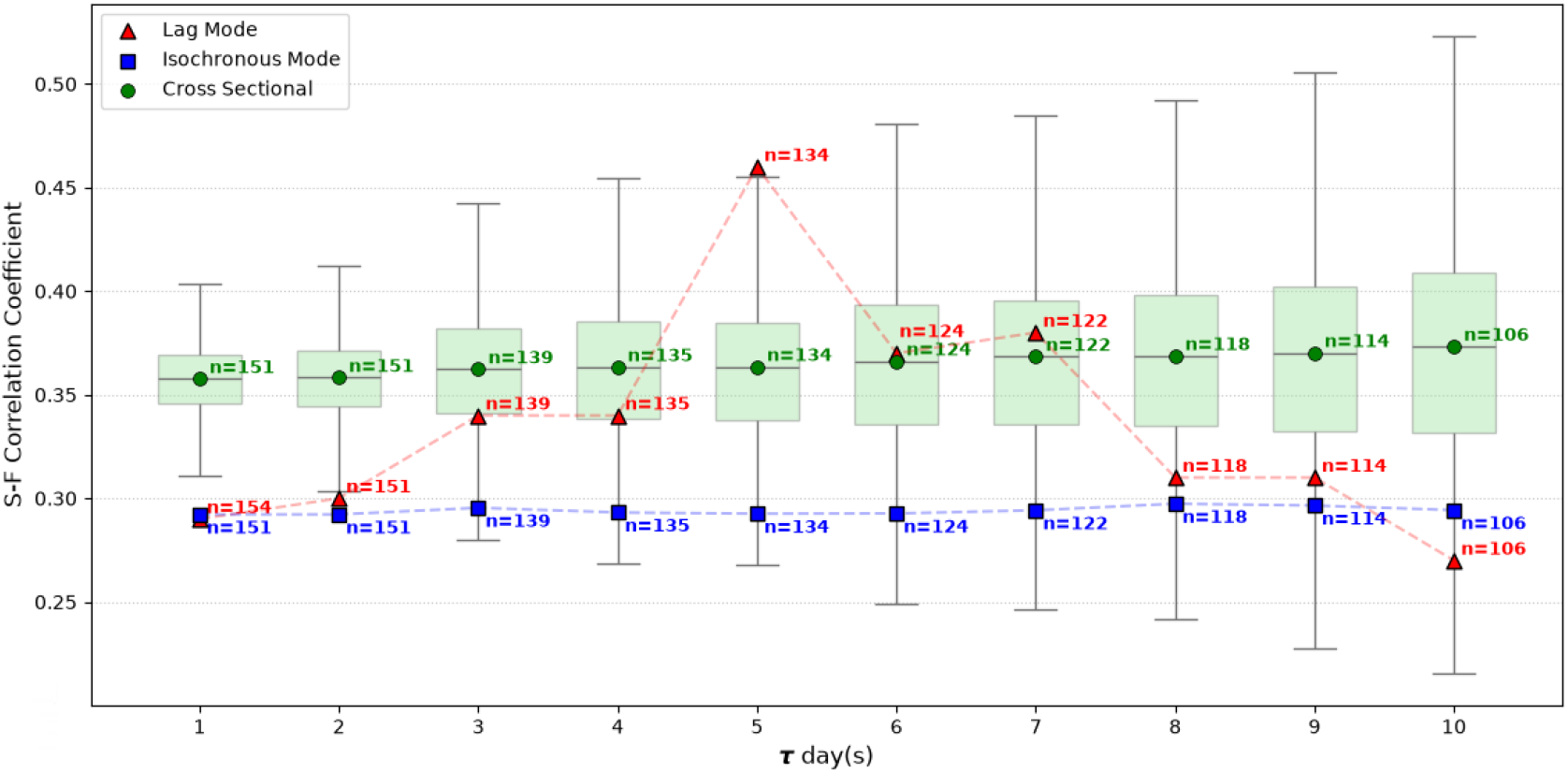
Comparing the S-F correlation values in lag (r =1-10 days) and isochronous (r =0) modes and in individuals in cross-sectional data. Red triangles: lag mode. Blue squares: isochronous mode. Green circles and boxplots: The S-F correlations computed on individuals from original cross-sectional data. The number of subject pairs in isochronous mode and the number of individuals in cross-sectional data were mostly matched to the number of subject pairs in lag mode.

As the age gap between paired subjects increases from 1 to 5 days, the correlation between the functional features of the older subject and the structural features of the younger subject progressively increases. This increase in S-F correlation provides strong evidence that structural maturation exerts delayed effects on the functional development of perisylvian regions. Notably, the maximum S-F correlation value (r=0.46) is observed at an age gap of 5 days, suggesting an estimation of the delay of functional maturation relative to structural maturation in perisylvian regions during the first weeks after birth. Beyond an age gap of 5 days, the S–F correlation exhibits a descending trend.

Furthermore, the correlations obtained using the proposed longitudinal pairing framework exceed those computed from individual structural and functional features in the cross-sectional dataset--even after resampling to match the number of subject pairs--at age gaps of 5 to 7 days. Notably, at the 5-day age gap, the correlation derived from the longitudinal pairing framework surpasses the maximum correlation observed within the distribution of resampled cross-sectional data. This finding provides further evidence that functional maturation in the perisylvian region follows structural maturation with an approximate delay of 5 days.

## DISCUSSION

Characterizing early brain development is crucial for predicting long-term neurobehavioral disabilities, particularly in preterms^30^. Efforts to characterize brain maturation were focused on creating dynamic structural^31–33^ and functional^30,34^ atlases, whereas the integration of structural and functional information into a joint S-F atlas, offering a more comprehensive representation of the developing brain, remains lacking. A major challenge in constructing joint atlases is identifying reliable biomarkers that accurately characterize S-F coupling. Among the increasingly sophisticated multimodal biomarkers proposed to characterize S-F coupling, only a few, relying on myelination as a structural feature or on structural-functional covariance, have reported consistent estimates of this relationship during early development^5– 7,11,12,35,36^. The generally weak and often statistically insignificant association between structural and functional features may, at least in part, stem from the implicit assumption that structural and functional maturation occur synchronously. This assumption may, however, underestimate the true relationship, because structural maturation and functional organization do not necessarily evolve simultaneously during early brain development. Our findings suggest incorporating a new temporal dimension to the current synchronous S-F correspondence, which offers a more meaningful characterization of neonatal brain development and improves the estimation of S-F coupling.

A key obstacle to incorporating temporal asynchrony into S-F relationships is the need for longitudinal neuroimaging data, particularly difficult to acquire during the neonatal period. To address this limitation, we proposed a longitudinal subject-pairing framework based on systematically acquired cross-sectional data. Rather than correlating the structural and functional features of the same individual, the proposed framework estimates asynchronous S-F correlation by associating the structural features of one subject with the functional features of another subject of different age. In this way, cross-sectional data are leveraged to approximate longitudinal developmental trajectories, partially overcoming the limited availability of true longitudinal datasets.

By relaxing the conventional assumption of synchronous structural and functional maturation, the proposed framework generalizes existing approaches for modeling neonatal S-F relationships. Beyond improving the early S-F maturation coupling, this strategy may provide a generalizable framework for studying developmental processes in scenarios where longitudinal data are difficult or impossible to acquire, while sufficiently dense cross-sectional datasets are available. Examples include investigations of normal brain development^37^, disease progression in neurological and neurodevelopmental disorders^10,38,39^, and treatment-related brain reorganization, where densely sampled cross-sectional cohorts may serve as a practical surrogate for longitudinal observations.

The proposed framework can provide a meaningful approximation of longitudinal S-F relationships from cross-sectional data only if a number of methodological issues are adequately addressed. The first challenge is to identify pairs of subjects within a cohort that exhibit a specified age difference while remaining mutually comparable in terms of their structural and functional brain features. To address this challenge, we formulated a cost function that exploits the inherent structural and functional similarities, as well as the corresponding inter-subject variability, between candidate subject pairs. Minimization of the cost function would yield an optimal set of subject pairs for a given age gap. The use of inter-subject anatomical similarity has previously been exploited in the construction of neonatal brain atlases^30–34^. However, the key contribution of the proposed framework lies in simultaneously incorporating both structural and functional similarities to determine the optimal correspondence between subjects. To our knowledge, this is the first study to integrate these complementary sources of information for subject pairing and subsequently estimate S-F coupling by correlating the structural features of a subject with the functional features of an older subject.

The proposed cost function for optimal subject pairings consists of a weighted sum of distance terms each representing a specific correspondence criterion. Determining the weighting coefficients is another important challenge. In this study, we performed an exhaustive search within the parameter space to identify the combination of paired subjects that achieve the highest S-F correlation. The exhaustive search within the parameter space inevitably increases the computational cost of the optimization process, particularly when applied to large cross-sectional neuroimaging cohorts. This trade-off between computational complexity and optimization accuracy should therefore be considered when scaling the proposed framework to substantially larger datasets. Nevertheless, for the neonatal cohort investigated in the present study, the computational burden associated with the exhaustive search remained entirely manageable and did not constitute a practical limitation.

The existence of a temporal lag between structural and functional maturation during early brain development is well established in neuroscience^8,14,40^. The delayed functional maturation of the perisylvian region relative to its structural development, as previously reported in^11,41^, was selected as a case study to evaluate the proposed framework. Using myelination index and fALFF as structural and functional features, respectively, extracted from the dHCP cross-sectional cohort, the proposed framework not only reproduced this established developmental phenomenon but also, to the best of our knowledge, provides the first quantitative estimate of its temporal delay in term neonates. Quantifying such developmental delays from cross-sectional rather than longitudinal observations represents a significant methodological advance. Consequently, the proposed framework provides a new opportunity to investigate the temporal dynamics of asynchronous S-F development of the brain with substantially greater detail than has previously been possible.

The functional maturation delay of 5 days is specific to the age range of 37–45 weeks PMA, as investigated in the present study. A factor that is likely to influence this temporal offset is the rate of development in structural and functional features. During periods of rapid brain maturation, the delay between structural and functional changes may be relatively short, whereas slower developmental periods may be associated with a longer temporal offset. Although this hypothesis remains to be systematically investigated, the proposed framework paves the way for examining age-dependent asynchrony in structure-function maturation across different developmental stages. Recent indirect evidence supports this interpretation, demonstrating that synchronous S-F coupling progressively transitions toward decoupling within auditory and somatosensory regions^42^, particularly following approximately 41 weeks PMA^43^. This observation is consistent with the possibility that the temporal offset between structural and functional maturation increases beyond the age range examined in the present study. Such a developmental shift is also biologically plausible, as structural processes, including axonal growth and maturation, may continue to strengthen structural connectivity without immediately producing corresponding changes in functional brain activity. Indeed, substantial increases in cortical activation have been reported to emerge only several months after birth, when widespread functional maturation becomes established^44^. Furthermore, structural and functional maturation are known to follow regionspecific developmental trajectories and maturation rates. It is therefore reasonable to expect that the temporal offset between structure and function may vary across different cortical regions. Investigating region-specific optimal delays represents an important direction for future work and may provide a more detailed understanding of the spatiotemporal organization of early brain development.

This study introduced a framework that leverages cross-sectional datasets^37,39,45^ by enabling longitudinal analyses through the pairing of subjects of different ages. This contribution may reduce the dependence on large-scale longitudinal acquisitions, which are often difficult, costly, and time-consuming in neonatal populations. Moreover, by providing quantitative estimates of developmental delays, the proposed framework may help optimize future longitudinal study designs by narrowing the temporal windows that require direct follow-up. It may also influence the design of future cross-sectional imaging protocols, including the selection of age sampling intervals and acquisition strategies that maximize their utility for reconstructing developmental trajectories.

More broadly, the proposed framework extends conventional approaches for joint S-F brain analysis by explicitly incorporating temporal asynchrony into the estimation of S-F coupling. This extension enables the investigation of questions that cannot be adequately addressed under the conventional assumption of synchronous S-F development. Although demonstrated here in the context of neonatal brain development, the underlying methodology may also be applicable to other domains in which structural and functional changes evolve over time but longitudinal observations are difficult to obtain, such as studies of brain aging^46^ and the progression of neurodegenerative disorders^47^, including dementia.

The proposed framework represents an initial step toward reconstructing joint S-F developmental trajectories from cross-sectional neonatal data. Future methodological developments may build upon this framework by integrating advanced trajectory inference techniques, including mathematical formulations based on optimal transport and related trajectory reconstruction methods that have recently demonstrated considerable success in reconstructing temporal progression from cross-sectional observations in cancer evolution and single-cell genomics^22,23^. Adapting such approaches to developmental neuroimaging may provide a powerful framework for modeling continuous S-F brain maturation.

The present study focused on the widely accepted developmental hypothesis that structural maturation provides the biological substrate upon which functional organization gradually emerges. Future studies could use this framework in lead mode to investigate the reciprocal relationship between structure and function, particularly in contexts such as experience-dependent brain plasticity, where functional activity is known to induce structural reorganization^17,48^.

## RESOURCE AVAILABILITY

### Lead contact

Requests for further information and resources should be directed to and will be fulfilled by the lead contact, Hamid Abrishami Moghaddam.

### Data and code availability

- This paper analyzes the second release of the dHCP dataset. The dHCP dataset is publicly available data, accessible at [https://www.developingconnectome.org/data-release/second-datarelease/].
- All original code and extracted features required to reanalyze the data reported in this paper are available from the lead contact upon request.

## ACKNOWLEDGMENTS

This work is partially supported by the Iran National Science Foundation (INSF) under the grant number 40409501.

## AUTHOR CONTRIBUTIONS

Conceptualization, H. Abrishami Moghaddam; Methodology, H. Abrishami Moghaddam, R. Namiranian, M. Sadeghi; Investigation, M. Sadeghi and R. Namiranian; Original draft, R. Namiranian and M. Sadeghi; Writing – review & editing, H. Abrishami Moghaddam and R. Namiranian; Funding acquisition, H. Abrishami Moghaddam; Resources, H. Abrishami Moghaddam; Supervision, H. Abrishami Moghaddam.

